# Examining the molecular mechanisms contributing to the success of an invasive species across different ecosystems

**DOI:** 10.1101/2020.03.23.003434

**Authors:** Sarah K. Lamar, Ian Beddows, Charlyn G. Partridge

## Abstract

Invasive species provide an opportune system to investigate how populations respond to new environments. Baby’s breath (*Gypsophila paniculata*) was introduced to North America in the 1800’s and has since spread throughout the United States and western Canada. We used an RNA-seq approach to explore how molecular processes contribute to the success of invasive populations with similar genetic backgrounds across distinct habitats. Transcription profiles were constructed from seedlings collected from a sand dune ecosystem in Petoskey, MI (PSMI) and a sagebrush ecosystem in Chelan, WA (CHWA). We assessed differential gene expression and identified SNPs within differentially expressed genes. We identified 1,146 differentially expressed transcripts across all sampled tissues between the two populations. GO processes enriched in PSMI were associated with nutrient starvation, while enriched processes in CHWA were associated with abiotic stress. Only 7.4% of the differentially expressed transcripts contained SNPs differing in allele frequencies of at least 0.5 between populations. Common garden studies found the two populations differed in germination rate and seedling emergence success. Our results suggest the success of *G. paniculata* in these two environments is likely due to plasticity in specific molecular processes responding to different environmental conditions, although some genetic divergence may be contributing to these differences.

## INTRODUCTION

The ability of invasive species to invade, adapt, and thrive in novel ecosystems has long been a focus of ecological research. Coined the “paradox of invasions”, examining how invasive populations respond to novel environmental stressors after an assumed reduction in population size during introduction has become an entire field of scientific inquiry [1–3]. However, this paradox has been called into question as research shows that while many invasive populations may undergo a reduction in demographic and/or effective population size after an invasion event, this is not always linked with a subsequent reduction in genetic diversity [1,4]. Additionally, differences between the total genetic diversity of a population and the adaptive variation of a population can be large [5,6]. For these reasons, using total genetic diversity as a measure of invasive potential can be complex and potentially misleading. Instead, a better approach may be to examine how invasive species functionally respond to novel environments and assess how specific molecular processes may be contributing to invasive success [3,7,8].

Local adaptive evolution and phenotypic plasticity represent two strategies for coping with novel environmental stressors, although they are not mutually exclusive [7,8]. Phenotypic plasticity can be adaptive, maladaptive, or neutral, and can occur independently or in conjunction with shifts in allele frequencies that also alter mean trait values [9,10]. When phenotypic plasticity is adaptive, the population’s trait value moves closer to the new environment’s optimum. This can allow populations to persist through the sudden application of strong directional selection that often accompanies an introduction, particularly a founder event, without the more time consuming process of having to wait for fortuitous mutations to arise [9,11–13]. Over time, if there are population distributional changes in allele frequencies associated with fitness, then the invasive population will have on average a phenotype that is more fit in its current range than it would be in other environments, including the native range. Regardless of the mechanism, these shifts in fitness-related traits are the difference between persistence and perishing for an introduced population [7,14,15].

In the study of invasive species, the ability to examine molecular processes associated with phenotypically plastic responses (e.g., through environmentally driven gene expression differences) and those indicative of local adaptive evolution (e.g., through changes in allele frequencies) is often limited by the relative lack of background genetic data available, particularly for non-model species [16]. Examining these two processes can be further complicated when traditional methods used to assess local adaptation, such as reciprocal translocation experiments, bring up ethical concerns since moving invasive populations to new locations may increase their potential spread [17]. This concern may be especially true for highly prolific invasive species. However, with the development of technologies like RNA-seq, which allows for the assembly of transcriptomes *de novo*, gene expression and sequence data have become more widely available for non-model systems [3,16,18]. RNA-seq derived gene expression data can be used to answer questions related to how different environments influence changes in gene expression, which can help address how plastic these responses may be [8,19,20]. In addition, because RNA-seq also produces sequence data, we can assess allele frequency differences for genes that are differentially expressed, which can give initial insight into population divergence and potential processes driving local adaptive evolution [21]. Thus, the combination of expression and sequence data produced from RNA-seq methods can allow researchers to estimate the prevalence of plasticity in response to novel environmental stressors and begin to address questions about how invasive species adapt to their introduced environments [3,8].

In this study we take advantage of RNA-seq technology to examine changes in different molecular processes that may allow invasive populations with similar genetic backgrounds to establish across different ecosystems. The system we are using to explore this question is invasive populations of baby’s breath (*Gypsophila paniculata* L.; Caryophyllaceae), which inhabits different regions of the continental U.S. and Canada. *Gypsophila paniculata* is a perennial forb native to Eurasia. It is a thought to be a long-lived herbaceous perennial (at least 7 years, CGP pers. obs.), although the full life span has not been assessed, and flowers are not produced until the second or third year of growing [22]. As is characteristic of most members of the genus *Gypsophila*, it thrives in environments with dry, well-draining, calcareous soils with warm summers and cool winters [23]. However, it has one of the largest geographic distributions of the genus, stretching from eastern Europe to North China [23,24]. Originally introduced into North America in the late 1800’s for use in the floral industry [22,25], *G. paniculata* quickly spread and can now be found growing in diverse ecosystems across North America, often outcompeting and crowding out the native species [26,27]. While relatively little is known about the history of invasive baby’s breath populations in the United States, a recent population genetic analysis using 14 microsatellite markers identified at least two distinct population clusters, with one of these clusters including populations that span from the upper portion of Michigan’s lower peninsula to the eastern side of the Cascade Mountains [28]. The environments that these populations occur in range from quartz sand dunes in Michigan, disturbed roadsides in Minnesota, prairies in North Dakota, and sagebrush steppes in eastern Washington. While these populations may share a similar genetic background, understanding how they are responding to different environments will help shed light on how this invasive is able to thrive across distinct habitats.

For this study, we examined differential gene expression and identified single nucleotide polymorphisms (SNPs) within differentially expressed genes from two *G. paniculata* populations within the same genetic cluster that inhabit divergent ecosystems: (1) the coastal sand dunes in Petoskey, Michigan and (2) sagebrush steppe regions around Chelan, WA. These two habitats were chosen because they represent ecologically distinct ecosystems, with divergent environmental characteristics (see results). In addition, we conducted a common garden growth trial to examine differences in germination rates, seedling emergence success, and above- and below-ground tissue allocation between these two populations. We predict that the populations will differ in gene expression patterns and that those differences will be reflective of the environment in which they inhabit. Given that baby’s breath established in these environments approximately 100 years ago [28], we also predict that this should be enough time to see divergence in allele frequencies for genes that are important to these distinct habitats. This will allow us to identify potential targets of local adaptive evolution for future testing. Finally, we hypothesize that different environmental conditions (i.e., growing degree day, precipitation, and nutrient availability (see results)) between these two habitats has likely led to differences in growth responses. Therefore, we predict that these populations will differ in certain phenotypic traits, such as germination rate, seedling emergence success, and above- and below-ground tissue allocation, when grown in a common garden environment. Thus, the overall goal of this work was to examine how *G. paniculata* populations that have shared genetic backgrounds but differ in their invaded habitats (i.e., sand dunes in Petoskey, Michigan, and sagebrush steppe in Chelan, Washington) are responding to these different environments and to explore how different molecular processes are contributing to their success as an invasive species.

## MATERIALS AND METHODS

### Study Site Characterization

Petoskey, Michigan (PSMI) is a state park located along Lake Michigan’s primary successional quartz-sand dune system. Vegetation is sparse and is chiefly comprised of *Ammophila breviligulata* (dune grass), *Silene vulgaris* (bladder campion), *Juniperus horizontalis* (creeping juniper), *J. communis* (common juniper), and *Cirsium pitcheri* (Pitcher’s thistle) (Figure 1a-b). Chelan, Washington (CHWA) is a disturbed habitat situated on slopes surrounding Lake Chelan and dominated by sagebrush (*Artemisia* spp.) (Figure 1a & c). Average climate data for these two locations were collected from stations operated by the National Oceanic and Atmospheric Organization (NOAA) in Petoskey, MI and Entiat, WA (near Chelan, WA) and is summarized in Table 1.

**Table 1.** Location and climate data for sampling sites, taken from National Oceanic and Atmospheric Organization (NOAA) weather stations in Petoskey, MI and Entiat, WA (near Chelan, WA).

|  | Station ID | GPS<br>Coordinates | Elevation<br>(m) | 2017 Mean<br>Temp. (°C) | 2018 Mean<br>Temp. (°C) | 2017<br>Precipitation<br>(cm) | 2018<br>Precipitation<br>(cm) | 2017<br>GDD | 2018<br>GDD |
| --- | --- | --- | --- | --- | --- | --- | --- | --- | --- |
| Entiat Fish<br>Hatchery (WA) | USC00452563 | 47.6983°, -<br>120.3228° | 313 | 10.33 | 12.22 | 37.95 | 27.81 | 3013 | 3050 |
| Petoskey (MI) | USC00206507 | 45.3725°, -<br>84.9766° | 182.6 | 7.66 | 7.17 | 109.75 | 88.62 | 2130 | 2178 |

\*GDD (Growing Degree Day) as defined by the National Weather Service; an approximation of crop maturity

**Figure 1.**
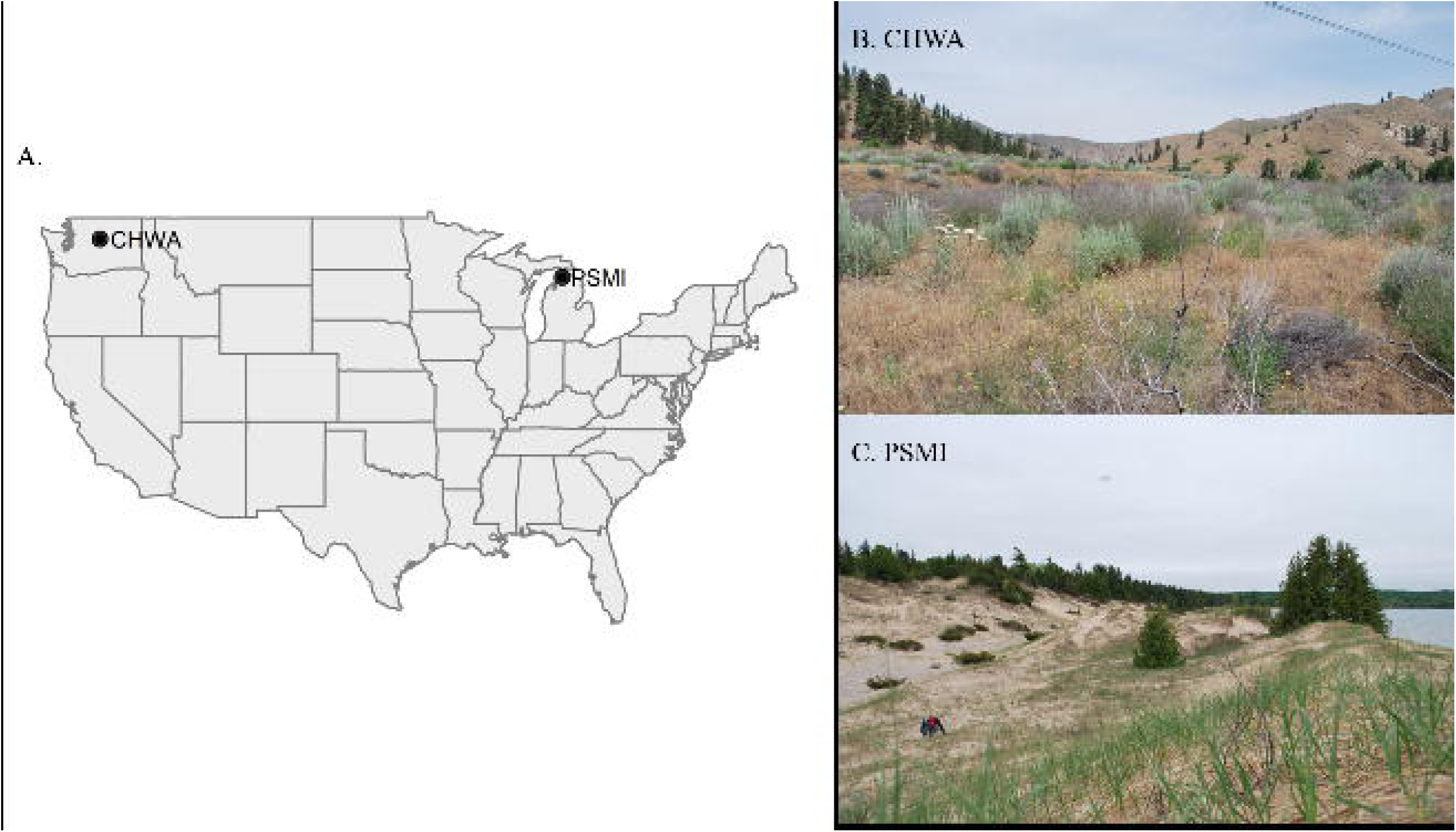
(A) Map identifying sample locations for *Gypsophila paniculata* populations used in this study. (B) Petoskey, Michigan (PSMI) study site, and (C) Chelan, Washington (CHWA) study site.

### Soil Analysis

In June of 2018, we collected soil samples from PSMI (47.7421°N 120.2177°W) and CHWA (45.4037°N 84.9121°W) (Figure 1a-c). In PSMI, we collected soil from 10 cm, 50 cm, and 1 m, while in CHWA, we collected soil from 10 cm, 25 cm, and 50 cm depths. Sampling locations differed in collection depths due to soil characteristics in CHWA that made deeper collection impossible (large boulders, hard soil). At both locations, we collected two sets of soil samples from all depths. We stored samples in airtight plastic bags and maintained them at 4°C until analysis.

We sent soil samples collected from all depths at PSMI and CHWA to A&L Great Lakes Laboratories (Fort Wayne, IN) for nutrient analysis. Samples were tested for: organic matter (%), phosphorus (P), potassium (K), magnesium (Mg), calcium (Ca), soil pH, total nitrogen (N), cation exchange capacity (CEC), and percent cation saturation of K, Mg, and Ca. At the laboratories, samples were dried overnight at 40°C before being crushed and filtered through a 2 mm sieve. The following methods were then used for each analysis: organic matter content (loss on ignition at 360°C), pH (pH meter), phosphorus, potassium, magnesium, and calcium content (Mehlich III Extraction and inductively coupled plasma mass spectrometry). Total nitrogen was determined using the Dumas method (thermal conductance). Results of nutrient testing were analyzed using a principal component analysis (PCA) in the statistical program R v3.6.0 [29].

### RNA Extraction

Along with soil samples, we collected 16 *G. paniculata* seedlings from CHWA (June 8, 2018) and 15 seedlings from PSMI (June 1, 2018). We then dissected seedlings into three tissue types (root, stem, and leaf), placed tissue in RNAlater™ (Thermo Fisher Scientific, Waltham, MA), and flash-froze them in an ethanol and dry ice bath. Samples were kept on dry ice for transport and maintained at −80°C until RNA extractions were performed.

We extracted total RNA from frozen tissue using a standard TRIzol® (Thermo Fisher Scientific) extraction protocol (https://assets.thermofisher.com/TFS-Assets/LSG/manuals/trizol_reagent.pdf). We resuspended the extracted RNA pellet in DNase/RNase free water. The samples were then treated with DNase to remove any residual DNA using a DNA-Free Kit (Invitrogen, Carlsbad, CA). We assessed RNA quality with a Bioanalyzer 2100 (Agilent Technologies, Santa Clara, CA) and NanoDrop™ 2000 (Thermo Fisher Scientific). RNA Integrity Number (RIN) values for individuals used in this study ranged from 6.1-8.3. However, because both chloroplast and mitochondrial rRNA can artificially deflate RIN values in plant leaf tissue, we deemed these values to be sufficient for further analysis based upon visualization of the 18S and 28S fragment peaks [30]. This resulted in high quality total RNA from 10 PSMI leaf, 10 PSMI stem, 10 PSMI root, 10 CHWA leaf, 9 CHWA stem, and 10 CHWA root samples. Finally, we submitted the total RNA samples to the Van Andel Research Institute for cDNA library construction and sequencing.

### cDNA Library Construction and Sequencing

Prior to sequencing, all samples were treated with a Ribo-Zero rRNA Removal Kit (Illumina, San Diego, CA). cDNA libraries were constructed using the Collibri Stranded Library Prep Kit (Thermo Fisher Scientific) before being sequenced on a NovaSeq 6000 (Illumina) using S1 and S2 flow cells. Sequencing was performed using a 2 × 100 bp paired-end read format and produced approximately 60 million reads per sample, with 94% of reads having a Q-score >30 (Table S2).

### Transcriptome Assembly

Prior to transcriptome assembly, read quality was assessed using FastQC (https://www.bioinformatics.babraham.ac.uk/projects/fastqc/). Adapters and bases with a quality score less than 20 were first trimmed from the raw reads using Trim Galore (https://www.bioinformatics.babraham.ac.uk/projects/trim_galore/). Next, rRNAs were identified using SortMeRNA (mean rRNA percent content of 5.31%) [31]. A reference transcriptome was then assembled *de novo* using non-rRNA reads from all samples and Trinity v2.8.2 [32,33] with a normalized max read coverage of 100, a minimum k-mer coverage of 10, and k-mer size set to 32. The assembled transcriptome was annotated using Trinotate v3.1.1. Trinotate was given open reading frames (ORFs) predicted from TransDecoder and transcript homology (blastx and blastp) to the manually curated UniProt database [34]. The final assembly consisted of 223,810 putative genes and 474,313 putative transcripts (N50 = 3,121) from the 59 samples.

### Differential Expression

To quantify transcript expression, reads were mapped back to the assembly using bowtie and quantified using the RSEM method as implemented in Trinity. Counts were generated for genes and transcripts. We then tested for differential gene expression using edgeR v3.22.5 in R v3.5.2 [29,35]. First, however, the count data was filtered and only transcripts with greater than 10 counts in at least 10 samples were included. Following filtering, 111,042 genes (49.61%) and 188,108 transcripts (39.66%) remained. Considering tissue type, 127,591 transcripts remained in the data from 20 root samples (26.90%), 125,261 transcripts remained in the 19 stem tissue samples (26.41%), and 112,499 transcripts remained in the 20 leaf tissue samples (23.72%). For differential expression testing, the data were stratified by tissue and filtered transcripts were then fit to the negative binomial (NB) model and tested using the quasi-likelihood F test with TMM (trimmed mean of M values) normalization. To be considered significantly differentially expressed, transcripts needed to have an adjusted p-value (BH method [36]) below 0.05 and a log2 fold change greater than 2.

For transcripts that were differentially expressed, we identified Gene Ontology (GO) biological processes that were either over- or under-represented using the PANTHER classification system v14.1, where transcripts were assessed against the *Arabidopsis thaliana* database (http://pantherdb.org/webservices/go/overrep.jsp). In addition, for those transcripts that were differentially expressed across all three tissues, we converted the UniProt IDs of the transcripts to GO biological process IDs using the online database bioDBnet (https://biodbnet-abcc.ncifcrf.gov/db/db2db.php), and used the metacoder package v0.3.3 [37] in R v3.6.0 to construct heat trees to visualize the relationship of our differentially expressed transcripts across GO biological process hierarchies.

### Single Nucleotide Polymorphism (SNP) Variant Calling

We used the HaplotypeCaller tool from GATK4 to identify potential SNPs that were present in transcripts that were differentially expressed between populations [38,39]. The bowtie mapped files were used to jointly genotype all 59 samples simultaneously with a minimum base quality and mapping quality of 30. Variant data was visualized using the vcfR package v1.8.0 [40].

We identified variants associated with non-synonymous SNPs, synonymous SNPs, 5’ and 3’ UTR SNPs, 5’ and 3’ UTR indels, frame-shift and in-frame indels, premature or changes in stop codons and changes in start codons, and calculated population diversity estimates for all SNP types. The effect prediction was done using custom scripts (which can be found in the Dryad repository) and the Transdecoder predicted annotation in conjunction with the base change. We set a hard filter for the SNPs so that only those with QD scores > 2, MQ scores > 50, SOR scores < 3, and Read Post Rank Sums between −5 and 3 passed. We then calculated the allele frequencies for each SNP within PSMI and CHWA. For the subsequent evaluation, we focused on SNPs that had potential functional effects (i.e., they were not listed as ‘synonymous’ or ‘unclassified’), were in transcripts differentially expressed between PSMI and CHWA across all three tissues, and that exhibited differences in SNP allele frequencies between the populations by at least 0.5. We used the R package metacoder v0.3.3 [37] to visualize the GO biological process hierarchies associated with transcripts containing these SNPs.

### Common Garden Trials

Finally, to examine whether environmental differences between these two locations has led to different growth responses, we conducted common garden trials to examine differences in germination rate (functionally defined as radicle emergence [41]), seedling emergence success (defined as successful cotyledon emergence from the soil), and the ratio of above- and below-ground tissue allocation between the populations.

#### Germination Trial

On August 11, 2018 we returned to our sample sites in CHWA and PSMI and collected seeds from 20 plants per location. This date was chosen because it was previously determined that this collection time can yield over 90% seed germination for *G. paniculata* collected from Empire, MI [42]. To collect seeds, we manually broke seed pods off and placed them inside paper envelopes in bags half-filled with silica beads. We stored bags in the dark at 20 to 23°C until the germination trial began one month later.

We counted one hundred seeds from twenty plants per population and placed them in a petri dish lined with wet filter paper (n = 2,000 seeds per population). We established a control dish using 100 seeds from the ‘Early Snowball’ commercial cultivar *(G. paniculata*) sold by W. Atlee Burpee & Co in 2018, known to have germination percentages in excess of 90%. Incubators had a 12:12h dark:light photoperiod and growth chamber conditions were set at 20°C with 114 μmol m^−2^ s^−1^ photosynthetically active radiation from fluorescent light bulbs. Each day we randomized petri dish locations within the incubator to avoid bias in temperature or light regimes. We conducted this study for fourteen days, at which point there had been no germination in any dish for two days. The same individual checked all seeds (n=4,100) daily within the same three-hour time window to minimize bias for germination. Once a seed had germinated, we removed it from the dish (method adapted from [42]).

Using the statistical program R v3.6.0, we fit the data to a nonparametric Kaplan-Meier time-to-event curve [29,43]. We then compared germination patterns between CHWA and PSMI using a pairwise log-rank test [43]. To test for homogeneity within localities, we again conducted a log-rank test. Finally, to investigate the presence of family effects (i.e., differences among seeds from different parental plants), we ran a series of pairwise log-rank tests with a Holm correction for multiple comparisons [43]. For all analyses in this study, we set the alpha level to 0.05.

#### Growth Trials

To examine population differences on seedling emergence success and above- and below-ground tissue allocation, we planted 6 seeds collected from 20 individual plants per population (n = 120 per population, n = 240 total). All seeds were planted on the same day to a standardized depth of 5 mm in a sand/potting soil mixture. Greenhouse conditions were set at 7:17 h dark:light photoperiod. Relative humidity and temperature settings during the day were 55% and 21°C while nighttime conditions were 60% and 15.5°C. Each day we watered plants until the soil appeared fully wet and we randomized plant position to prevent bias in temperature, light, or water regime. At the end of the seven-week trial period, we carefully removed plants from the soil and measured the length of tissue above and below the caudex using a caliper.

To compare the proportion of seedlings that successfully emerged between the populations, we ran a two-sided proportion test in the R statistical program v3.6.0 [29]. We analyzed differences in the ratio of above- and below-ground tissue between populations for seedlings that successfully emerged and examined the presence or absence of family effects using a completely randomized design with subsampling ANOVA in SAS v9.4 [44].

## RESULTS

### Habitat Characterization

Climate data collected from NOAA monitoring stations revealed differences in mean temperature, precipitation, and growing degree day (GDD) between our two sampling locations. CHWA had a 3°C and 5°C higher mean temperature in 2017 and 2018 than PSMI, while PSMI had greater rainfall in both 2017 (109.8 cm vs 38 cm) and 2018 (88.6 cm vs 27.8 cm) (Table 1). CHWA had a greater number of GDD in both 2017 (3,013 vs. 2,130) and 2018 (3,050 vs. 2,178) (Table 1). Soils collected from CHWA were characterized by higher levels of total nitrogen, phosphorus, magnesium, and potassium. In contrast, soils from PSMI had a higher pH and more available calcium (Figure 2, Table S1).

**Figure 2.**
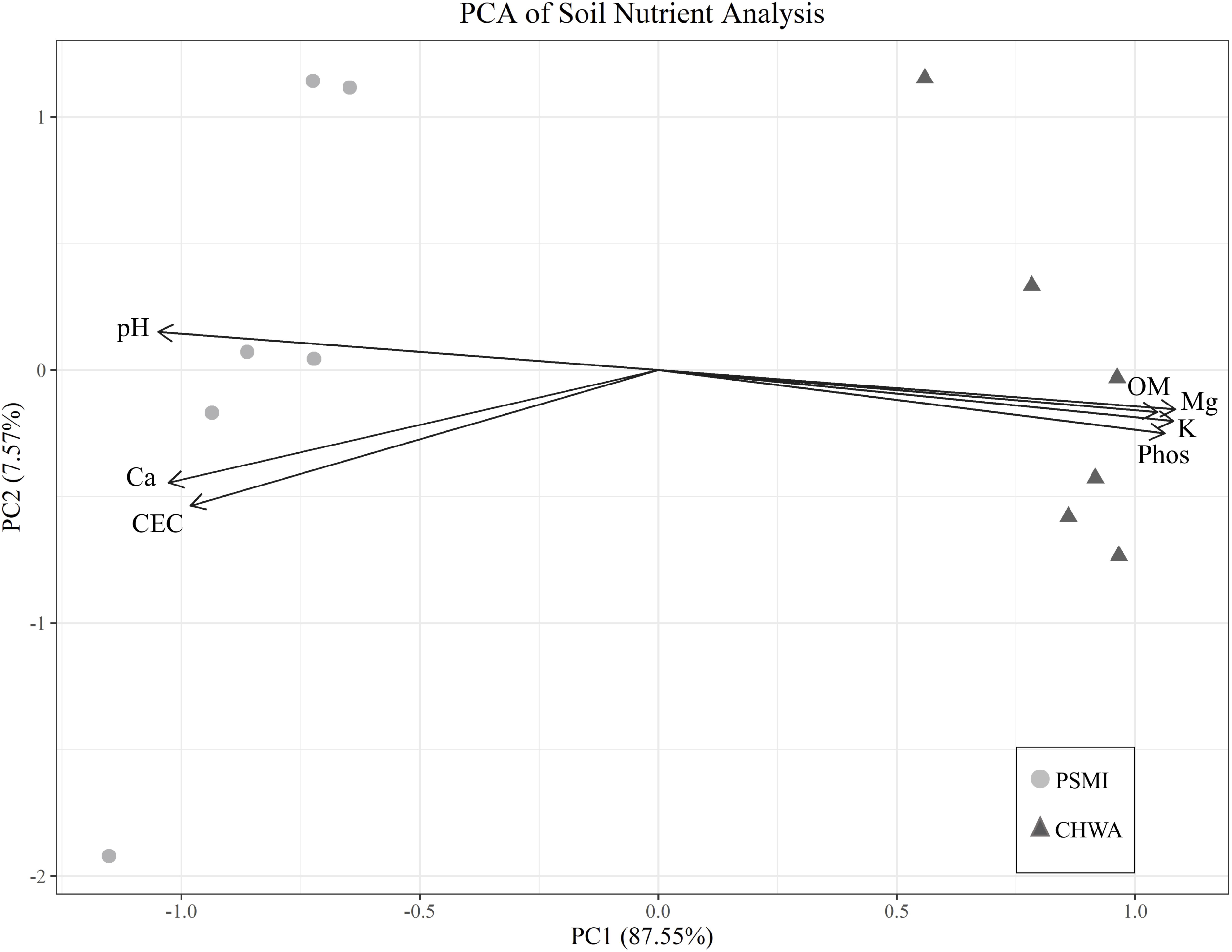
Principal component analysis (PCA) of soil nutrient data for sampling locations in Petoskey, Michigan (PSMI) and Chelan, WA (CHWA). Ca = calcium, CEC = cation exchange capacity, K = potassium, Mg = magnesium, OM = organic matter, and Phos = phosphorus

### Differential Gene Expression

Across all three tissue types, there were 1,146 transcripts that were differentially expressed between the PSMI and CHWA populations (Figure 3a, Table S3), with the majority of the differences in expression being driven by sampling location and tissue type (Figure 3b). Root tissue contained the highest number of differentially expressed transcripts between the two populations (8,135 transcripts, Table S4), followed by leaf tissue (5,666 transcripts, Table S5) and stem tissue (5,376 transcripts) (Figure 3a, Table S6).

**Figure 3.**
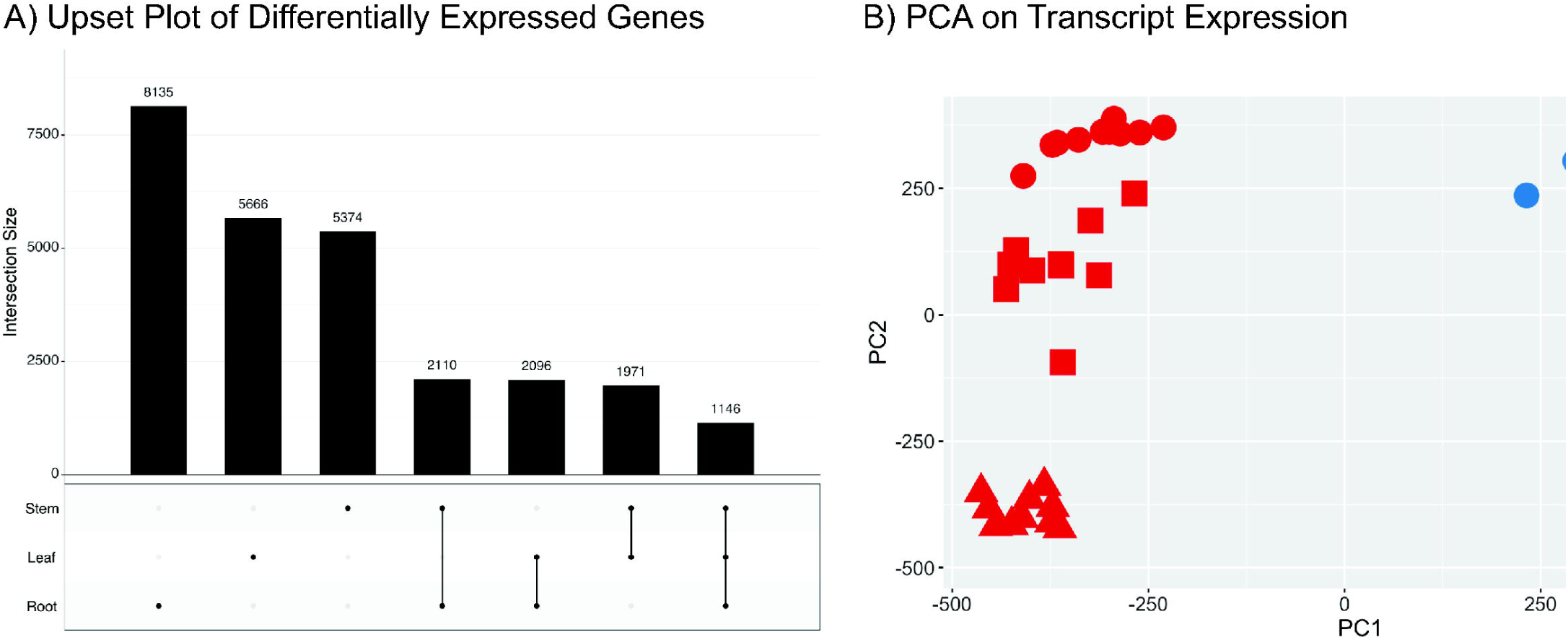
Differential transcript expression among groups. (A) Upset intersection plot visualizing the number of differentially expressed transcripts in *Gypsophila paniculata* growing in Chelan, Washington (CHWA) and Petoskey, Michigan (PSMI) broken down by tissue type (root, stem, and leaf tissue). The bar chart represents the number of differentially expressed transcripts in each tissue type, while the below-chart matrix illustrates the tissue type(s) being considered in the analysis. Dark grey connected circles indicate transcripts that are differently expressed across multiple tissues. B) PCA plot associated with transcript expression profiles.

### Enriched GO Processes Between Populations

#### Enriched GO Processes in CHWA

GO biological processes that were enriched with transcripts displaying higher expression in CHWA relative to PSMI across all three tissue types were primarily associated with different stress responses (Table 2). These included response to reactive hydrogen species (GO:0000302), cellular response to unfolded proteins (GO:0034620), protein import into the peroxisome matrix (GO:0016560), response to heat (GO:0034605), response to water deprivation (GO:0009414), and response to abscisic acid (GO:0009737). Many of the stress response related GO processes included a number of heat shock protein genes that displayed higher expression in CHWA across the three tissues (Table S3).

**Table 2:**
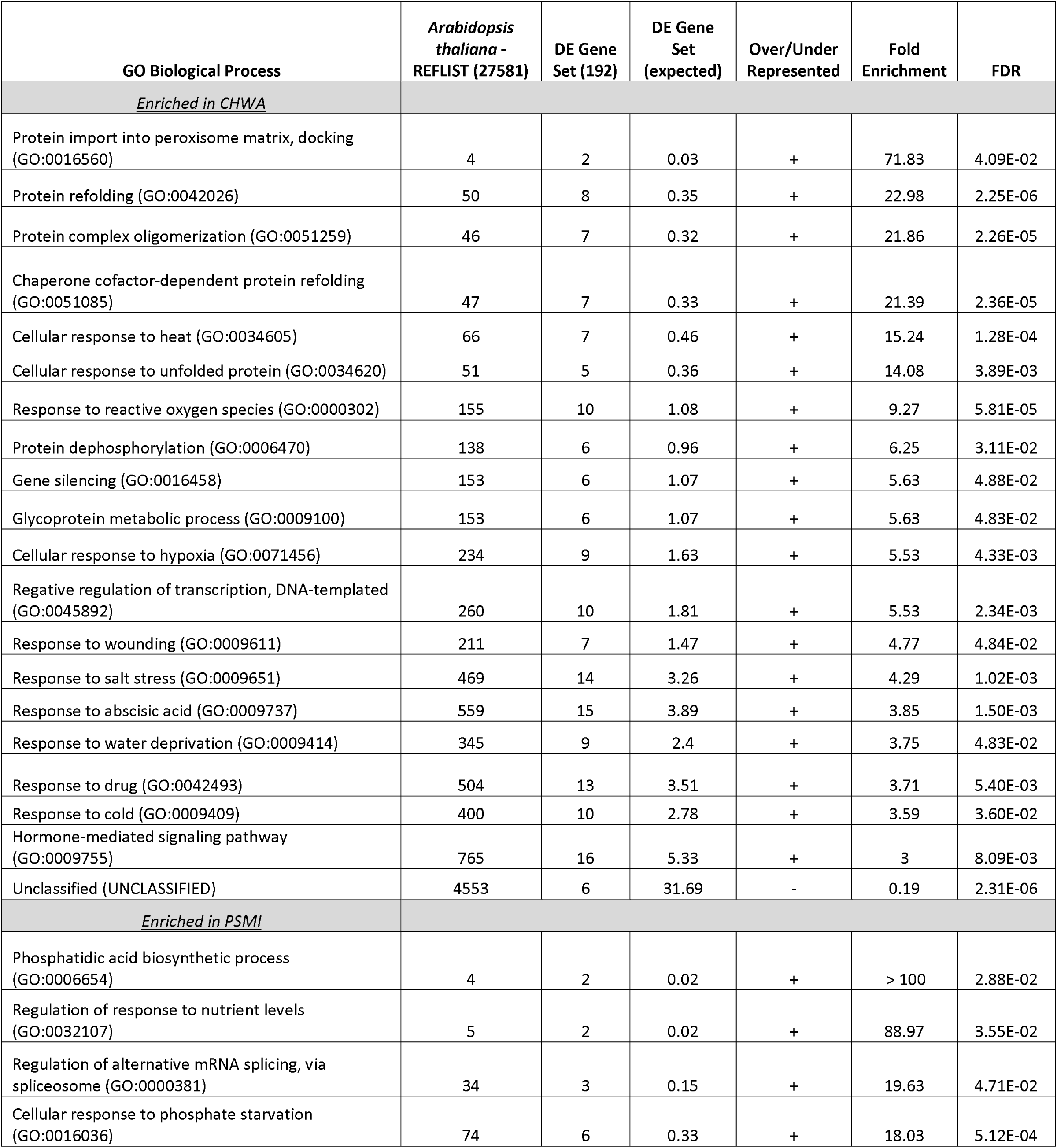
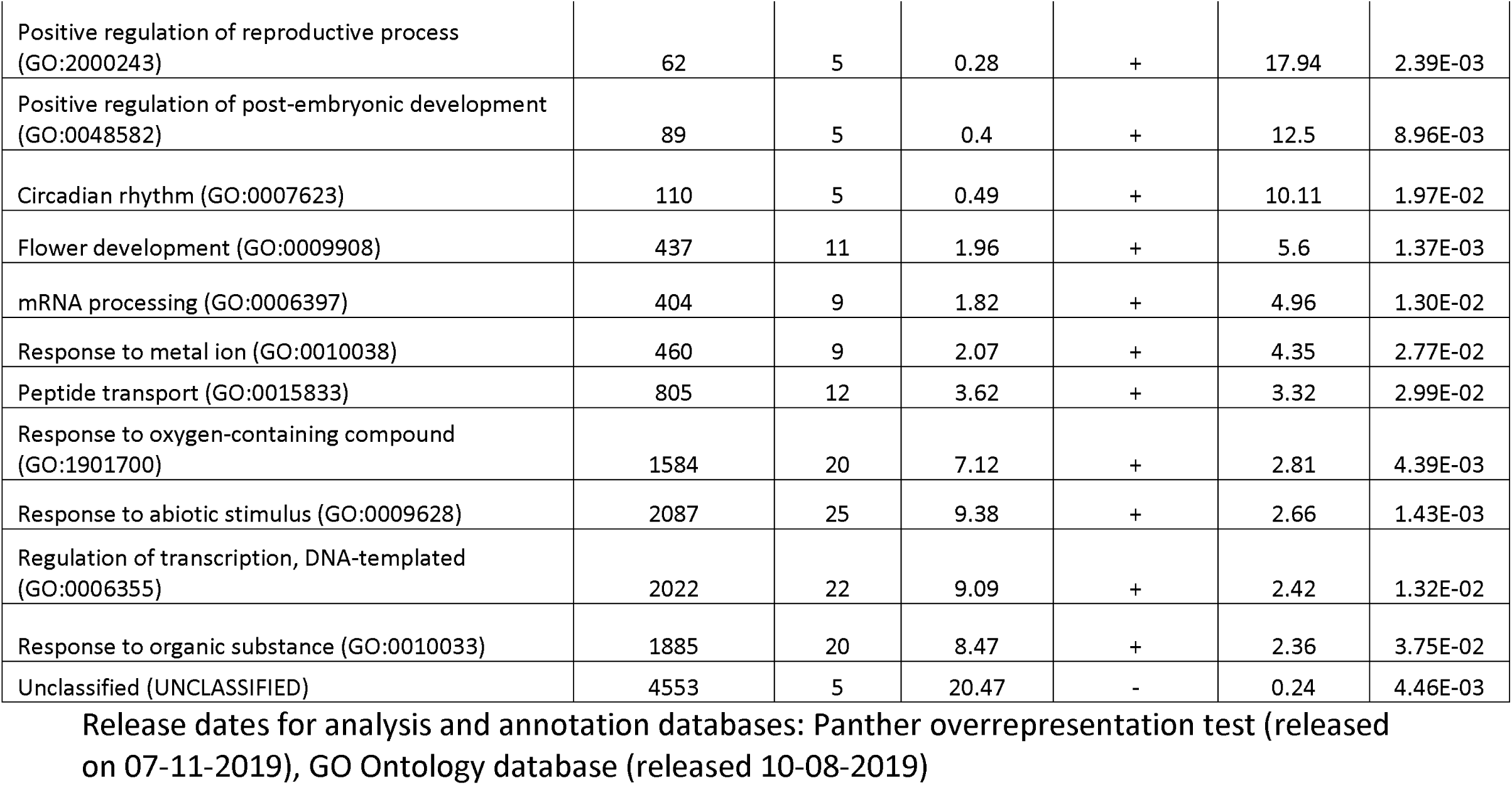
GO biological processes enriched across all three tissues (root, stem, and leaf) for Petoskey, MI (PSMI) and Chelan, WA (CHWA) populations. DE: Differentially expressed, FDR = False Discovery Rate.

#### Enriched GO Processes in PSMI

For the PSMI population, GO terms that were enriched with transcripts that showed significantly higher expression across all three tissues were associated with nutrient response, development, and transcriptome processes (Table 2). These included regulation of response to nutrient levels (GO:0032107), cellular response to phosphate starvation (GO:0016036), phosphatidic acid biosynthesis process (GO:0006654), response to metal ion (GO0010038): circadian rhythm (GO:0007623), flower development (GO:0009908), regulation of alternative mRNA splicing via spliceosome (GO:0000381), and regulation of DNA-templated transcription (GO:0006355). Transcripts associated with multiple GO terms related to nutrient processes included phospholipase D zeta 2 (*PLPZ2*), transcription factor HRS1 (*HRS1*), and SPX domain containing protein 3 (*SPX3*). Some of the circadian rhythm and flower development associated transcripts included Adagio protein 3 (*ADO3*), protein GIGANTEA (*GIGAN*), and lysine-specific demethylase JMJ30 (*JMJ30*). A comparison of GO biological processes hierarchies associated with transcripts differentially expressed between the two populations can be visualized in Figure 4.

**Figure 4.**
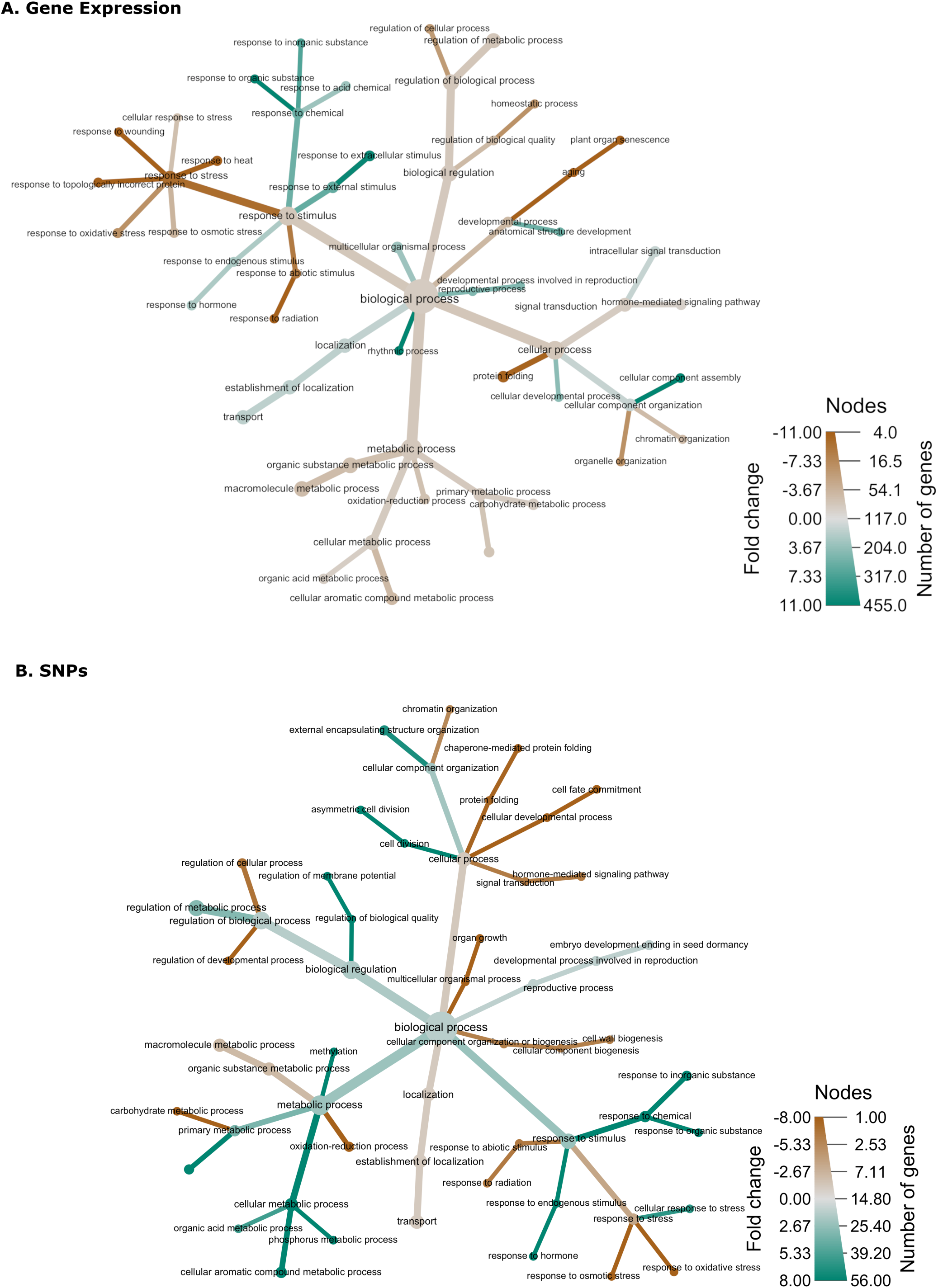
Heat trees displaying (A) GO biological processes that are enriched with transcripts with significant differential expression between each population, and (B) GO biological processes that are represented by transcripts differentially expressed between the two populations and contain SNPs that differ in allele frequency by at least 0.5. The size of each node is representative of the number of transcripts assigned to each GO term. The color of each branch represents increased expression, with green displaying higher expression in Petoskey, Michigan (PSMI) and brown displaying higher expression in Chelan, Washington, (CHWA).

### Tissue Specific Gene Expression Patterns

#### Root Tissue

When directly comparing transcript expression between root tissue from CHWA and PSMI, 63% (5,131 transcripts) were more highly expressed in CHWA, while 37% (3,004 transcripts) displayed higher expression in PSMI. Enriched GO terms from the CHWA population specifically associated with root tissue may be involved in defense and/or stress responses and included cellular response to salicylic acid stimulus (GO:0071446) and regulation of plant-type hypersensitive response (GO:0010363). In addition, processes associated with COPI coating of golgi vesicles (GO:0048205) and xyloglucan metabolic processes (GO:0010411) were specifically enriched in CHWA root tissue (Table S7). For the PSMI population, GO terms specifically associated with root tissue included cellular response to nitrogen starvation (GO:0006995), nitrate assimilation (GO:0042128), and organophosphate metabolic processes (GO:0019637) (Table S8).

#### Stem Tissue

There were 5,374 differentially expressed transcripts in stem tissue collected from CHWA and PSMI (Figure 3a). Of those, 2,421 transcripts (45%) displayed higher expression in CHWA while 2,953 transcripts (55%) were more highly expressed in PSMI. For the CHWA stem tissue, specific GO processes included response to sucrose (GO:0009744), regulation of response to DNA damage stimulus (GO:2001020), and telomere maintenance in response to DNA damage (GO:0043247) (Table S9). Processes that were specific to the PSMI stem tissue included phosphoenolpyruvate transport (GO:0015714) and systemic acquired resistance (GO:0009627) (Table S10).

#### Leaf Tissue

Of the 5,666 transcripts that were differentially expressed between leaf tissue from CHWA and PSMI (Figure 3a), 58% (3,286 transcripts) displayed higher expression in CHWA compared to the 42% (2,380 transcripts) that showed relatively higher expression in PSMI. Some of the enriched GO terms that were specific to leaf tissue from the CHWA population included fatty acid beta-oxidation (GO:0006635), and positive regulation of salicylic acid mediated signaling pathway (GO:0080151) (Table S11). The enriched GO terms that were specific to PSMI leaf tissue included, vitamin biosynthesis process (GO:0009110), long-day photoperiodism and flowering (GO:0048574), and response to UV-A (GO:0070141) (Table S12).

### Comparison of Gene Expression and SNP GO Biological Processes

Of the transcripts that were differentially expressed between CHWA and PSMI across all three tissues, 85 (7.4%) of those transcripts contained potentially functional SNPs, which displayed allele frequencies that differed between the two populations by at least 0.5 (Table S13). Enrichment analysis did not identify any GO processes that were statistically enriched for these 85 transcripts; although, GO biological terms associated with these transcripts can be viewed in Figure 4b.

### Germination Trial

Results of a log-rank test comparing time-to-germination curves for each locality indicated strong statistical differences between seeds collected from PSMI and CHWA, with seeds from CHWA germinating more quickly (p < 2.0 × 10^−16^) (Figure 5). While there was a difference in germination curves, both localities reached 90% germination by the end of the germination trial. Log-rank tests looking at homogeneity within groups found strong statistical support for variation among time-to-germination curves for seeds from different parent plants for both populations (both p < 2.0 × 10^−16^), suggesting potential family effects.

**Figure 5.**
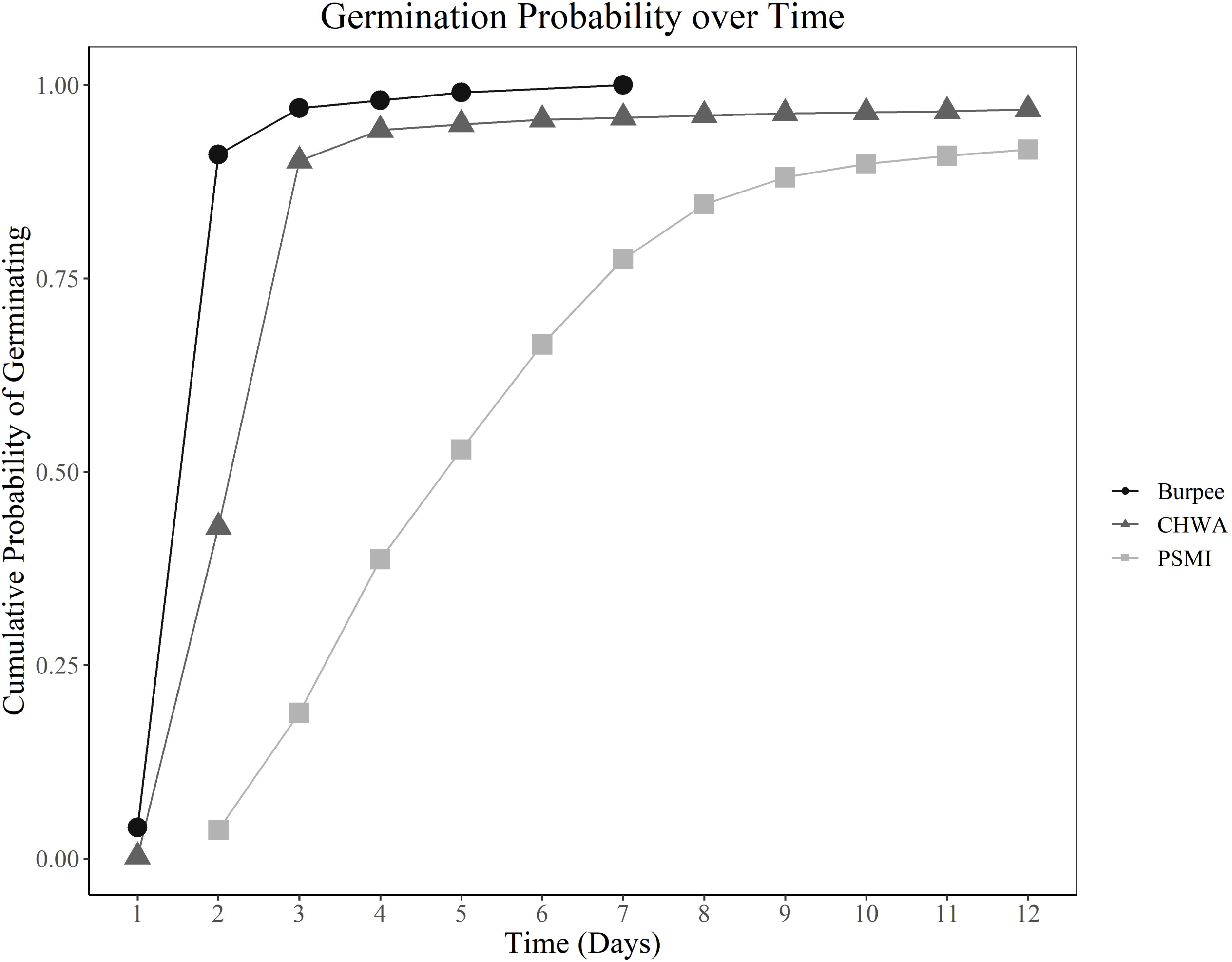
Germination curves for *Gypsophila paniculata* seeds collected from Petoskey, Michigan (PSMI, n = 2,000) and Chelan, Washington (CHWA, n = 2,000) on August 11, 2018 and incubated for 12 days. Burpee commercial cultivar seeds (n = 100) known to have germination success in excess of 90% were used for an experimental control.

### Growth Trial

A two-sided proportion test indicated a significant difference in the total number of seedlings that emerged between seeds collected from CHWA and PSMI, with CHWA seedlings emerging more often than PSMI (p<0.0002) (Figure 6a). When excluding plants that did not emerge, ANOVA results indicated no significant difference in the ratio of above- and below-ground tissue allocation between populations (p=0.605) (Figure 6b). However, there were significant family effects in above- and below ground tissue allocation (p=0.0301) (Figure S1).

**Figure 6.**
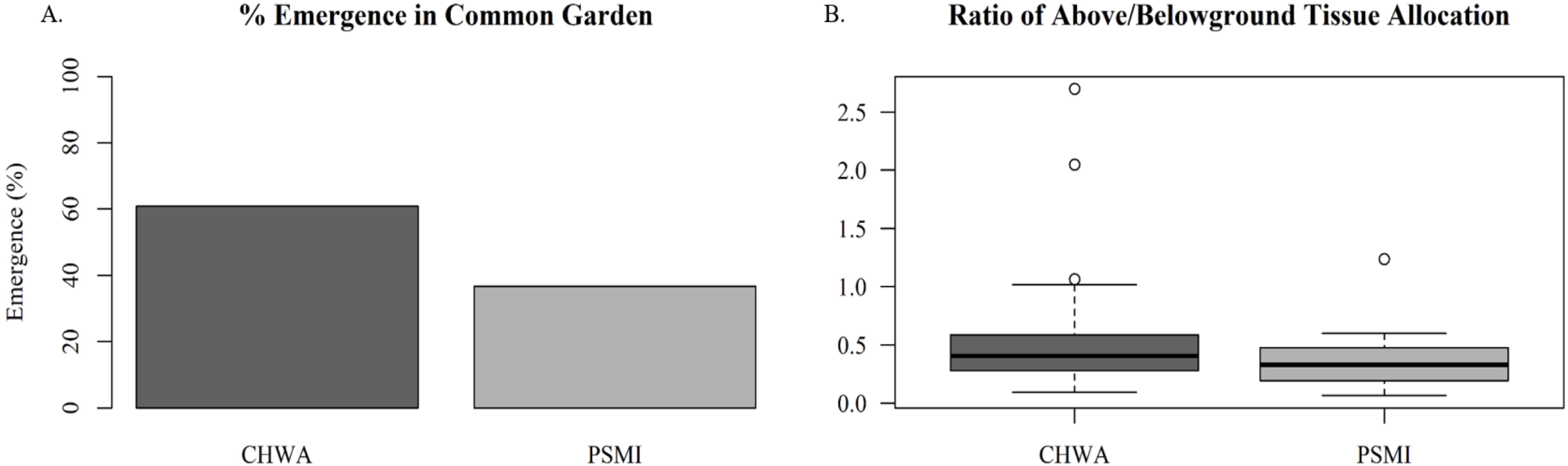
Results of a common garden growth trial of *Gypsophila paniculata* plants conducted for seven weeks (n=120 per population). (A) Seedling emergence per sampling location, (B) Ratio of above: below-ground tissue allocation per sampling location. Location codes: Chelan, Washington (CHWA); Petoskey, Michigan (PSMI).

## DISCUSSION

The primary drivers that allow invasive species to adapt to novel environments over relatively short periods of evolutionary time is a process not yet fully understood. To better understand these mechanisms, we investigated two populations of *G. paniculata* growing at opposite ends of the species’ introduced range in the United States. Based on herbarium records, *G. paniculata* populations have been established in the sand dunes of Petoskey, MI since the early 1910’s and in the sagebrush steppes of Washington since the 1930’s [28]. This has likely provided sufficient time for these populations to begin adapting to these divergent ecosystems. However, genetic analysis of North American *G. paniculata* at neutral microsatellite loci show that these two populations belong to the same genetic cluster, suggesting a shared history [28]. Using RNA-seq data (which gives orders of magnitude more informative data than microsatellites), we found that there were a number of transcripts differentially expressed between these populations and that many of these genes were involved in processes directly related to their different environments, particularly those associated with abiotic stress response in CHWA and nutrient starvation in PSMI. Of the genes that were differentially expressed across all three tissues, only 7.4% contained potential SNPs that differed in frequency by at least 0.5 between the populations. In addition, while we identified differences in germination rates and seedling emergence success between the two populations in a common garden experiment, we did not observe differences in above- and below-ground tissue allocation as we initially predicted. From these data, we suggest that the success of invasive *G. paniculata* across these distinct ecosystems is likely the result of plasticity in molecular processes responding to these different environmental conditions, although some genetic divergence over the past 100 years may also be contributing to these differences.

### Stress Response in CHWA

The sagebrush ecosystem of the eastern Cascade Mountains is characterized by a semi-arid, temperate environment with a drought-resistant plant community [45]. The environmental data obtained from our sampling regions suggests that the CHWA population experiences less precipitation and higher temperatures than *G. paniculata* growing in PSMI. As such, many of the enriched GO processes with higher expression in the CHWA population were related to a suite of stress responses indicative of abiotic stress. Some of these included response to abscisic acid (ABA), response to reactive oxygen species, response to heat, response to salt stress, response to water deprivation, and response to topologically incorrect folded proteins (Table 2, Figure 4a). During abiotic stress, many of these processes interact with one another to help maintain cellular homeostasis [46,47]. In our data, transcripts that were associated with protein folding GO processes mainly corresponded to heat-shock proteins (Hsps). While Hsps are most notably involved in protein stability during heat stress, they can also respond when plants experience osmotic, cold, or oxidative stress [48–51]. Hsps can also interact with ABA, often considered a ‘plant stress hormone’ because it can be induced by multiple abiotic stressors [52,53]. *Arabidopsis* mutants that are deficient in ABA do less well under drought or osmotic stress conditions than those with sufficient ABA [47]. Under heat and drought stress, increased production of ABA can lead to higher levels of hydrogen peroxide and result in oxidative stress. But, this effect can be mediated as increased oxidative stress triggers synthesis of *Hsp70*, which up-regulates antioxidant enzymes that control reactive oxygen species and protects against oxidative injury [54,55]. Thus, enrichment of genes involved in these interacting processes suggests CHWA populations are under higher levels of abiotic stress, particularly heat and drought stress, compared to PSMI populations and these data provide insight into the molecular response to these stressors.

When examining leaf, root, stem tissue from CHWA seedlings separately, additional GO processes related to stress responses were observed. ‘Response to salicylic acid’ was enriched in both the leaf and root tissue. Salicylic acid (SA) is a phytohormone that is involved in immunity and defense response to pathogens [56,57]. It also plays an important role in plant’s response to abiotic stress, including metal, salinity, ozone, UV-B radiation, temperature, and drought stress [58]. For example, in *Mitragyna speciose*, application of SA led to increased expression of chaperone proteins and heat shock proteins when plants were in drought conditions [58,59]. As previously stated, the arid environment of the sagebrush ecosystem is likely to result in higher drought stress, and increased expression of genes associated with SA pathways may be an additional mediating factor allowing invasive *G. paniculata* to thrive in this system.

While a number of genes involved in abiotic stress response showed higher expression in CHWA, the majority of these genes did not have SNPs with divergent allele frequencies between the two populations, suggesting that some of this response is likely due to plasticity. However, a few genes involved in different stress responses and chaperon-mediated protein folding processes did have SNPs that differed in allele frequency by at least 0.5 or greater. One of the genes involved in oxidative stress was caffeoylshikimate esterase (*CSE*). CSE is an important enzyme in the synthesis of lignin, a major component of the cell wall [60]. Plants with mutations in the *CSE* gene display increased sensitivity to hydrogen peroxide and oxidative stress, which were enriched in our GO analysis [61]. In addition, another transcript that displayed divergent allele frequencies was peptidyl-prolyl cis-trans isomerase (*FKBP62*), which is involved in chaperone-mediate protein folding. *FKBP62* interacts with the heat shock protein 90 (*HSP90.1*) complex to positively regulate thermotolerance in *Arabidopsis* [62]. Expression of this gene is induced in *Arabidopsis* during heat stress, and those that over-express this gene show higher survival at temperatures above 45° C after a 37 °C acclimation period [62]. This increased heat tolerance could be helpful in the warmer, arid climate of CHWA. Differences in allele frequencies between PSMI and CHWA associated with these genes suggest that there could be local adaptive evolution occurring due to different selection pressures associated with abiotic stress. However, additional work needs to be conducted to more thoroughly examine these distinct SNPs and fully assess their relationship to population divergence and local adaptive evolution.

### Nutrient Starvation in PSMI

The *G. paniculata* population in PSMI is located in the coastal sand dunes of northwest Michigan. This area is a primary-successional dune habitat where *G. paniculata* grows in the foredune region. The sand dune environment can present strong selection pressure on plants in the form of sand burial, limited soil moisture, and lack of nutrients [63]. One of the main limiting factors for seedling success in dune systems is nutrient deficiency, especially nitrogen, phosphorus, and potassium [64,65]. Our soil analysis show that PSMI soil contained low concentrations of organic matter, total nitrogen, phosphorus, and potassium, suggesting this is a very nutrient limited environment. In conjunction with these environmental differences, the GO enrichment analysis showed that ‘regulation of response to nutrient levels’ and ‘cellular response to phosphate starvation’ were both significantly enriched in PSMI in all three tissues compared to CHWA. In addition, there were a number of processes associated with nitrate regulation (nitrate assimilation and nitrogen cycle metabolic process) specifically enriched in the root tissue from PSMI. Some of the differentially expressed genes associated with these processes included phospholipase D zeta 2 (*PLPZ2*), transcription factor HRS1 (*HRS1*), and SPX domain containing protein 3 (*SPX3*). In *Arabidopsis thaliana, PLPZ2* can aid in phosphate recycling, and has been shown to be upregulated during phosphate starvation [66]. Additionally, *SPX3* helps regulate phosphate homeostasis [67,68], while *HRS1* is a major regulator of both nitrogen and phosphate starvation [69]. The increased expression of these genes may help *G. paniculata* survive in PSMI, where the limited levels of nitrate and phosphorus in the soil make this ecosystem a challenge for many plant species. However, these specific genes did not display SNPs that differed in frequency between our populations, suggesting that expression differences related to nutrient deprivation are environmentally driven, potentially epigenetically maintained, and/or are regulated by non-transcribed regions, and these differences exist in response to the low nitrogen and phosphorus environment experienced in the dune system.

When examining PSMI GO processes enriched with differentially expressed genes that contain SNPs differing in frequency between the two populations, the only nutrient-associated process was ‘phosphorus metabolic processes’. The gene involved in this process was CDP-diacylglycerol-glycerol-3-phosphate 3-phosphatidyltransferase 1 (*PGPS1*), which is involved in phosphatidylglycerol (PG) biosynthesis [70]. While this gene itself has not directly been associated with nutrient homeostasis, PG can be used as a phosphate reserve during phosphate starvation, and rapidly decreases in cells when phosphate is limited [71,72]. Thus, it is possible that the increase in *PGPS1* may be needed to maintain PG levels under these nutrient limited environments. However, further analysis needs to be performed to determine if the SNPs identified alter the function of this gene.

### Circadian Rhythm Expression in PSMI

There were also a number of enriched GO processes in PSMI related to different timing processes, including circadian rhythm and flowering-associated photoperiod. These two processes can be linked, with the circadian clock mechanisms that drive 24-hour cycles also significantly influencing plant phenology [73]. Ideally, circadian cycles should be optimized to match environmental parameters [74,75], and a disruption in circadian rhythm cycles can result in decreased fitness [76,77]. Given differences in both latitude and growing degree days between PSMI and CHWA, we would expect there to be differences in phenology between the populations, and this was evident during our collecting period. Even though we collected from both populations within one week of each other, and we tried to sample from both locations at the same time of day, some mature plants in CHWA were already budding, while mature plants in PSMI were still in the growth stage of their yearly life cycle. For most of the transcripts involved in these processes, there was not a corresponding SNP between the populations, suggesting these differences may be environmentally driven. However, a transcript associated with early flowering 3 protein (*ELF3*) displayed increased expression in the CHWA population and contained a SNP that differed in frequency between these populations. *ELF3* has been shown to modulate both flowering time and circadian rhythm [78], and interestingly, can also lead to increased salt tolerance (osmotic stress) in *Arabidopsis* [79]. These results suggest that environmental factors eliciting changes in timing and phenology may be helping to maintain these invasive populations.

### Phenotypic Comparisons: Germination and Growth Trials

To see what effects environmental factors might be having on different life history traits of our populations, we set up common garden growth trials. Different environmental factors can have varying selective pressures on germination rates, seedling emergence success, and above- and below-ground tissue allocation [80,81]. In our common garden experiments, we initially observed that seeds collected from CHWA germinated quicker and had higher seedling emergence success than those collected from PSMI. The better performance of the CHWA population could be due to release from the abiotic stress factors that were indicated by our gene expression data. Improved performance when a species is removed from an environment imposing abiotic stressors is a common hypothesis and is used as one explanation for the success of invasive species [82]. In this case, the high levels of drought and heat stress experienced in the sagebrush environment may enable the CHWA plants to have increased performance once these stressors have been removed. It is also possible that different selection pressures between the two environments could be leading to higher germination rates and seedling emergence success in CHWA. Lower precipitation in CHWA relative to PSMI could lead to seeds to be predisposed to germinate at the first instance of heavy watering. However, these differences could also be due to vegetative characteristics in the region. Specifically, *G. paniculata g*rowing in CHWA must compete against woody perennials (mainly *Artemisia sp.*) that are already established aboveground at the start of every growing season, while *G. paniculata* growing along the dune shore competes with grass species that sprout new leaves every year. Because survival is not dependent on merely whether the plants in CHWA can grow, but whether or not they can compete effectively, early germination could confer advantages in water limited environments, like a sagebrush steppe.

We saw no differences in above- and below-ground tissue allocation after seedling emergence between populations, suggesting there are no genetic differences between these populations in relation to these growth measures. We expected the nutrient limitation in PSMI to have an influence on the above- and below-ground tissue allocation of seedlings. In environments where nitrogen and phosphorus are the main limiting nutrients, root growth can be favored in seedlings relative to above-ground growth [83]. Additionally, nitrogen has been found to limit above-ground biomass in nutrient poor environments [84]. In contrast, shortage of Ca, which was present in higher quantities in PSMI than in CHWA, has been found to have little or no influence on above- and below-ground tissue allocation in lab experiments [83]. The lack of difference observed in root:shoot ratios in our plants could indicate that these factors do not influence tissue allocation resources in *G. paniculata* seedlings, or that these differences are not seen when *G. paniculata* is grown in a nutrient sufficient environment.

For our common garden trials, in addition to some of the population differences identified, we also observed significant family effects in germination rate, seedling emergence success, and above- and below-ground tissue allocation ratios, suggesting the potential for genetic effects. Variation in these traits are known to be driven, in part, by genetic factors in other plants. For example, for *Brassica oleracea*, heritability estimates of mean seed germination time and root:shoot length is approximately 14% and 12%, respectively. Looking to our gene expression data from our field collected seedlings, we did not observe differential expression of candidate genes proposed to be involved in germination timing (i.e. *AHG1, ANAC060, PDF1* [85]); however this could be due to the age of the seedlings upon collection. While the family effects we observed could be a function of genetic differences between seeds from different parental plants, these results can be significantly confounded by maternal effects. Seeds were collected from individual plants in their distinct environments, and different plants within these environments are likely allocating seed resources differently. In order to parse out genetic factors versus maternal effects associated with the environment, we would need to generate multiple generations within a common garden setting and examine gene expression differences once the maternal environment has been controlled. While this is something we hope to do in the future, it is beyond the scope of this current work. Regardless of the underlying cause, the data indicate that PSMI and CHWA populations display differences in life history traits that may be specific to the divergent environmental pressures present in these environments.

While this study is meant to serve as a first step in teasing apart how invasive *G. paniculata* populations are responding to different ecosystems, we acknowledge that there are additional variables that can alter the interpretation of these results. First, while we have data suggesting that PSMI and CHWA share similar genetic backgrounds [28], we do not know for certain the demographic history of these populations. Thus, the genetic differences that we are observing may be confounded by the past history of these populations prior to initial introduction to these areas. Secondly, in this study we only examined one population within a sand dune habitat and one population from a sagebrush habitat. Again, because demographic history can be a confounding factor, we cannot explicitly state that differences between these environments are solely driving the differences in gene expression patterns we observed or that SNP differences between these populations are not simply due to genetic drift. In the future, we plan to include more populations from each habitat, as well as additional prairie habitats, to explore this further. However, given the close relationship between the environmental characteristics of these habitats and the GO processes that were enriched within each population, we think that these processes are worthy of further evaluation of how molecular mechanisms may be driving the success of *G. paniculata* in these distinct ecosystems. Third, while RNA-seq analysis allowed us to examine SNPs in differentially expressed genes, there could also be genetic differences in non-transcribed regions that regulate gene expression between these populations. In these cases, some of the differential gene expression that we are observing could still be due to genetic differences between these populations, even though no SNPs were observed between the transcripts. To capture this information, further genetic analysis comparing these two populations would need to be conducted. Fourth, while we only identified a small number of differentially expressed genes with potentially functional SNPs that differed in allele frequency by 0.5 between the two populations, we acknowledge that this is a conservative cutoff and we have not considered the potential pleiotropic effects these genes may have on the different enriched processes. Additionally, further work needs to be conducted to identify any functional effects of these identified SNP differences and assess if they drive differences between populations. Finally, to fully assess local adaptation, more traditional approaches like reciprocal transplant experiments are needed. Although, given that *G. paniculata* is a prolific reproducer, transplanting more individuals into these sensitive habitats may bring significant ethical concerns. However, by identifying SNPs in differentially expressed genes that are divergent between these populations these data can provide an initial starting point to identify potential candidate genes that may be involved in adaptation to these novel habitats. Thus, regardless of these caveats, we feel that this work provides a good starting point toward identifying how different molecular processes influence *G. paniculata’s* success across these distinct ecosystems.

In conclusion, we found that *G. paniculata* seedlings from CHWA and PSMI displayed differential gene expression that was characteristic of the environment in which they were collected. In the nutrient limited sand dunes ecosystem, genes involved in responding to nutrients and phosphate starvation were upregulated. In the arid sagebrush ecosystem, genes involved in regulating responses to abiotic stress were upregulated. Given the small number of differentially expressed transcripts that contained divergent SNPs, we suggest that the majority of the expression differences associated with these enriched GO processes are likely driven by plastic responses to these different environments. Genetic divergence, however, cannot be completely dismissed given the differences in germination rates and seedling emergence success between the two populations in the common garden setting; although these seeds were collected from wild populations and maternal, environmental, and epigenetic variables could be contributing factors. Overall, this study reveals how variation in molecular processes can aid invasive species in adapting to a wide range of environmental conditions and stressors found in their introduced range.

## ACKNOWLEDGMENTS

We would like to thank Emma Rice and Hailee Leimbach-Maus for assistance during seed collection and with the germination study. We would also like to thank Jim McNair for help with statistical analysis for the common garden experiment, Marie Adams from the Van Andel Institute for library construction and sequencing, and Zachary Foster for help with Metacoder. We would also like to thank the Bureau of Land Management and the Michigan Department of Natural Resources for assistance with permitting and sample collection. Funding support was provided through Thermo Fisher Scientific, Grand Valley State University’s Presidential Research Grant, the Michigan Botanical Foundation, and the Environmental Protection Agency’s Great Lakes Restoration Initiative (C.G.P., Grant #00E01934).

## AUTHOR CONTRIBUTIONS

S.K.L. and C.G.P. conceptualized and designed this study. S.K.L. and C.G.P. carried out field collection and initial RNA extractions. I.B. performed the bioinformatics analysis, including transcriptome assembly, differential gene expression analysis, and SNP identification. C.G.P. performed the GO enrichment analysis. S.K.L. conducted the greenhouse and germination studies. S.K.L. and C.G.P. wrote the initial draft of the manuscript. All authors contributed to the final manuscript.

## DATA AVAILABILITY STATEMENT

All raw sequence reads associated with these data were deposited to the Sequence Read Archives (Bioproject accession #: PRJNA606240). Raw growth, germination data files, and R code for differential expression analysis and SNP identification are available on Dryad for review and will be made public once the manuscript is available (https://datadryad.org/stash/share/7XUtUP1t7wbBU6dd-hYM-FQss7XOknK2lf-HSCw9oSQ).

## Notes

doi:10.5061/dryad.v9s4mw6rq

### Competing Interest Statement

ThermoFisher funded the sequencing portion of the study in exchange for QA/QC data to assess their Collibri Stranded Library Prep Kits. ThermoFisher had no influence in terms of data reporting or data interpretation.

### Summary of Updates

Dryad link updated, Figure 2 revised. Manuscript updated, Figure 1 revised.

## REFERENCES

1. Dlugosch KM, Anderson SR, Braasch J, Cang FA, Gillette HD. 2015 The devil is in the details: Genetic variation in introduced populations and its contributions to invasion. Mol. Ecol. 24, 2095–2111. (doi:10.1111/mec.13183)

2. Sax DF, Brown JH. 2000 The paradox of invasion. Glob. Ecol. Biogeogr. 9, 363–371. (doi:10.3905/JOI.2010.19.1.032)

3. Sork VL. 2018 Genomic studies of local adaptation in natural plant populations. J. Hered. 109, 3–15. (doi:10.1093/jhered/esx091)

4. Frankham R. 2005 Resolving the genetic paradox in invasive species. Heredity (Edinb). 94, 385–385. (doi:10.1038/sj.hdy.6800634)

5. Leinonen T, O’Hara RB, Cano JM, Merilä J. 2008 Comparative studies of quantitative trait and neutral marker divergence: A meta-analysis. J. Evol. Biol. 21, 1–17. (doi:10.1111/j.1420-9101.2007.01445.x)

6. McKay JK, Latta RG. 2002 Adaptive population divergence: Markers, QTL and traits. Trends Ecol. Evol. 17, 285–291. (doi:10.1016/S0169-5347(02)02478-3)

7. Kawecki TJ, Ebert D. 2004 Conceptual issues in local adaptation. Ecol. Lett. 7, 1225–1241. (doi:10.1111/j.1461-0248.2004.00684.x)

8. Lande R. 2015 Evolution of phenotypic plasticity in colonizing species. Mol. Ecol. 24, 2038–2045. (doi:10.1111/mec.13037)

9. Ghalambor CK, McKay JK, Carroll SP, Reznick DN. 2007 Adaptive versus non-adaptive phenotypic plasticity and the potential for contemporary adaptation in new environments. Funct. Ecol. 21, 394–407. (doi:10.1111/j.1365-2435.2007.01283.x)

10. Van Kleunen M, Fischer M. 2005 Constraints on the evolution of adaptive phenotypic plasticity in plants. New Phytol. 166, 49–60. (doi:10.1111/j.1469-8137.2004.01296.x)

11. Conover DO, Schultz ET. 1995 Phenotypic similarity significance of countergradient variation. TRENDS Ecol. Evol. 10, 248–252. (doi:10.1016/S0169-5347(00)89081-3)

12. López-Maury L, Marguerat S, Bähler J. 2008 Tuning gene expression to changing environments: From rapid responses to evolutionary adaptation. Nat. Rev. Genet. 9, 583–593. (doi:10.1038/nrg2398)

13. Van Tienderen PH. 1997 Generalists, specialists, and the evolution of phenotypic plasticity in sympatric populations of distinct species. Evolution. 51, 1372–1380.. (doi:10.1111/j.1558-5646.1997.tb01460.x)

14. Joshi J et al. 2001 Local adaptation enhances performance of common plant species. Ecol. Lett. 4, 536–544. (doi:10.1046/j.1461-0248.2001.00262.x)

15. Richards CL, Bossdorf O, Muth NZ, Gurevitch J, Pigliucci M. 2006 Jack of all trades, master of some? On the role of phenotypic plasticity in plant invasions. Ecol. Lett. 9, 981–993. (doi:10.1111/j.1461-0248.2006.00950.x)

16. Ekblom R, Galindo J. 2011 Applications of next generation sequencing in molecular ecology of non-model organisms. Heredity. 107, 1–15. (doi:10.1038/hdy.2010.152)

17. Bunting D, Coleman RA. 2014 Ethical consideration in invasion ecology: A marine perspective. Ecol. Manag. Restor. 15, 64–70. (doi:10.1111/emr.12072)

18. Wang Z, Gerstein M, Snyder M. 2009 RNA-Seq: a revolutionary tool for transcriptomics. Nat. Rev. Genet. 10, 57–63. (doi:10.1038/nrg2484)

19. Des Marais DL, Hernandez KM, Juenger TE. 2013 Genotype-by-Environment Interaction and Plasticity: Exploring Genomic Responses of Plants to the Abiotic Environment. Annu. Rev. Ecol. Evol. Syst. 44, 5–29. (doi:10.1146/annurev-ecolsys-110512-135806)

20. Via S, Lande R. 2006 Genotype-Environment interaction and the evolution of phenotypic plasticity. Evolution. 39, 505. (doi:10.2307/2408649)

21. Costa V, Angelini C, De Feis I, Ciccodicola A. 2010 Uncovering the complexity of Transcriptomes with RNA-SEQ. J. Biomed. Biotechnol.207–247. (doi:10.1201/b16568)

22. Darwent AL, Coupland RT. 1966 Life History of Gypsophila paniculata. Weeds 14, 313. (doi:10.2307/4040974)

23. Barkoudah YI. 1962 A revision of Gypsophila, Bolanthus, Ankyropetalum and Phryna. Wentia 9, 1–203.

24. CABI. 2015 Gypsophila paniculata (baby’s breath). (online). See https://www.cabi.org/isc/datasheet/26266#toDistributionMaps (accessed on 19 March 2020).

25. Darwent AL. 1975 The biology of Canadian weeds: 14. Gypsophila paniculata L. Biol. Can. Weeds 55, 1049–1058. (doi:10.4141/cjps75-164)

26. Baskett CA, Emery SM, Rudgers JA. 2011 Pollinator visits to threatened species are restored following invasive plant removal. Int. J. Plant Sci. 172, 411–422. (doi:10.1086/658182)

27. Rice E. 2018 Assessment of invasive Gypsophila paniculata control methods in the northwest Michigan dunes. Masters Theses 888.

28. Lamar SK, Partridge CG. 2019 Old meets new: Combining herbarium databases with genetic methods to evaluate the invasion status of baby’s breath (Gypsophila paniculata) in North America. bioRxiv, 686691. (doi:10.1101/686691)

29. R Development Core Team. 2017 R: A language and environment for statistical computing.

30. Babu C. V. S, Gassmann M. 2016 Assessing integrity of plant RNA with the Agilent 2100 Bioanalyzer System. Agil. Appl. Note 5990-8850E.

31. Kopylova E, Noé L, Touzet H. 2012 SortMeRNA: Fast and accurate filtering of ribosomal RNAs in metatranscriptomic data. Bioinformatics 28, 3211–3217. (doi:10.1093/bioinformatics/bts611)

32. Grabherr MG et al. 2011 Full-length transcriptome assembly from RNA-Seq data without a reference genome. Nat. Biotechnol. 29, 644–52. (doi:10.1038/nbt.1883)

33. Haas BJ et al. 2013 De novo transcript sequence reconstruction from RNA-seq using the Trinity platform for reference generation and analysis. Nat. Protoc. 8, 1494–512. (doi:10.1038/nprot.2013.084)

34. Bryant DM et al. 2017 A Tissue-Mapped Axolotl De Novo Transcriptome Enables Identification of Limb Regeneration Factors. Cell Rep. 18, 762–776. (doi:10.1016/j.celrep.2016.12.063)

35. Robinson MD, McCarthy DJ, Smyth GK. 2009 edgeR: A Bioconductor package for differential expression analysis of digital gene expression data. Bioinformatics 26, 139–140. (doi:10.1093/bioinformatics/btp616)

36. Benjamini Y, Hochberg Y. 1995 Controlling the False Discovery Rate: A Practical and Powerful Approach to Multiple Testing. J. R. Stat. Soc. Ser. B 57, 289–300. (doi:10.1111/j.2517-6161.1995.tb02031.x)

37. Foster ZSL, Sharpton TJ, Grünwald NJ. 2017 Metacoder: An R package for visualization and manipulation of community taxonomic diversity data. PLoS Comput. Biol. 13, e1005404. (doi:10.1371/journal.pcbi.1005404)

38. McKenna A et al. 2010 The genome analysis toolkit: A MapReduce framework for analyzing next-generation DNA sequencing data. Genome Res. 20, 1297–1303. (doi:10.1101/gr.107524.110)

39. Depristo MA et al. 2011 A framework for variation discovery and genotyping using next-generation DNA sequencing data. Nat. Genet. 43, 491–501. (doi:10.1038/ng.806)

40. Knaus BJ, Grünwald NJ. 2017 vcfr: a package to manipulate and visualize variant call format data in R. Mol. Ecol. Resour. 17, 44–53. (doi:10.1111/1755-0998.12549)

41. Baskin JM, Baskin CC. 2001 Seeds: Ecology, Biogeography, and Evolution of Dormancy and Germination. 2nd edn. New York: Academic Press. (doi:10.1086/392994)

42. Rice EK, Martínez-Oquendo P, McNair JN. 2019 Phenology of seed maturation in babysbreath (Gypsophila paniculata) in northwest Michigan, USA, and its relation to glyphosate efficacy. Invasive Plant Sci. Manag. 12, 1–8. (doi:10.1017/inp.2019.21)

43. McNair JN, Sunkara A, Frobish D. 2012 How to analyse seed germination data using statistical time-to-event analysis: non-parametric and semi-parametric methods. Seed Sci. Res. 22, 77–95. (doi:10.1017/s0960258511000547)

44. SAS Institute Inc. 2013 SAS/ACCESS® 9.4.

45. Miller RF, Knick ST, Pyke DA, Meinke CW, Hanser SE, Wisdom MJ, Hild AL. 2011 Characteristics of Sagebrush Habitats and Limitations to Long-Term Conservation. Gt. sage-grouse Ecol. Conserv. a Landsc. species its habitats. (doi:10.1525/california/9780520267114.003.0011)

46. Shinozaki K, Yamaguchi-Shinozaki K. 2000 Molecular responses to dehydration and low temperature: differences and cross-talk between two stress signaling pathways. Curr. Opin. Plant Biol. 3, 217–223. (doi:10.1016/s1369-5266(00)80068-0)

47. Tuteja N. 2007 Abscisic acid and abiotic stress signaling. Plant Signal. Behav. 2, 135–138. (doi:10.4161/psb.2.3.4156)

48. Vierling E. 1991 The roles of heat shock proteins in plants. Annu. Rev. Plant Physiol. Plant Mol. Biol. 42, 579–620.

49. Boston RS, Viitanen P V., Vierling E. 1996 Molecular chaperones and protein folding in plants. In Post-Transcriptional Control of Gene Expression in Plants, pp. 191–222. Springer Netherlands. (doi:10.1007/978-94-009-0353-1_9)

50. Waters ER, Lee GJ, Vierling E. 1996 Evolution, structure and function of the small heat shock proteins in plants. J. Exp. Bot. 47, 325–338. (doi:10.1093/jxb/47.3.325)

51. Wang W, Vinocur B, Shoseyov O, Altman A. 2004 Role of plant heat-shock proteins and molecular chaperones in the abiotic stress response. Trends Plant Sci. 9, 244–252. (doi:10.1016/j.tplants.2004.03.006)

52. Swamy PM, Smith BN. 1999 Role of abscisic acid in plant stress tolerance. Curr. Sci. 76, 1220–1227.

53. Mahajan S, Tuteja N. 2005 Cold, salinity and drought stresses: An overview. Arch. Biochem. Biophys. 444, 139–158. (doi:10.1016/j.abb.2005.10.018)

54. Fauconneau B, Petegnief V, Sanfeliu C, Piriou A, Planas AM. 2002 Induction of heat shock proteins (HSPs) by sodium arsenite in cultured astrocytes and reduction of hydrogen peroxide-induced cell death. J. Neurochem. 83, 1338–1348. (doi:10.1046/j.1471-4159.2002.01230.x)

55. Hu X, Liu R, Li Y, Wang W, Tai F, Xue R, Li C. 2010 Heat shock protein 70 regulates the abscisic acid-induced antioxidant response of maize to combined drought and heat stress. Plant Growth Regul. 60, 225–235. (doi:10.1007/s10725-009-9436-2)

56. Dempsey DA, Shah J, Klessig DF. 1999 Salicylic Acid and Disease Resistance in Plants. CRC. Crit. Rev. Plant Sci. 18, 547–575. (doi:10.1080/07352689991309397)

57. Vlot AC, Dempsey DA, Klessig DF. 2009 Salicylic Acid, a Multifaceted Hormone to Combat Disease. Annu. Rev. Phytopathol. 47, 177–206. (doi:10.1146/annurev.phyto.050908.135202)

58. Khan MIR, Fatma M, Per TS, Anjum NA, Khan NA. 2015 Salicylic acid-induced abiotic stress tolerance and underlying mechanisms in plants. Front. Plant Sci. 6, 462. (doi:10.3389/fpls.2015.00462)

59. Jumali SS, Said IM, Ismail I, Zainal Z. 2011 Genes induced by high concentration of salicylic acid in Mitragyna speciosa. Aust. J. Crop Sci. 5, 296–303.

60. Vanholme R et al. 2013 Caffeoyl shikimate esterase (CSE) is an enzyme in the lignin biosynthetic pathway in arabidopsis. Science (80-.). 341, 1103–1106. (doi:10.1126/science.1241602)

61. Gao W, Li HY, Xiao S, Chye ML. 2010 Acyl-CoA-binding protein 2 binds lysophospholipase 2 and lysoPC to promote tolerance to cadmium-induced oxidative stress in transgenic Arabidopsis. Plant J. 62, 989–1003. (doi:10.1111/j.1365-313X.2010.04209.x)

62. Meiri D, Breiman A. 2009 Arabidopsis ROF1 (FKBP62) modulates thermotolerance by interacting with HSP90.1 and affecting the accumulation of HsfA2-regulated sHSPs. Plant J. 59, 387–399. (doi:10.1111/j.1365-313X.2009.03878.x)

63. Maun MA. 1994 Adaptations enhancing survival and establishment of seedlings on coastal dune systems. Vegetatio 111, 59–70. (doi:10.1007/BF00045577)

64. Willis AJ, Yemm EW. 1961 Braunton Burrows: Mineral Nutrient Status of the Dune Soils. J. Ecol. 49, 377. (doi:10.2307/2257270)

65. Hawke MA, Maun MA. 1988 Some aspects of nitrogen, phosphorus, and potassium nutrition of three colonizing beach species. Can. J. Bot. 66, 1490–1496. (doi:10.1139/b88-207)

66. Misson J et al. 2005 A genome-wide transcriptional analysis using Arabidopsis thaliana Affymetrix gene chips determined plant responses to phosphate deprivation. Proc. Natl. Acad. Sci. U. S. A. 102, 11934–11939. (doi:10.1073/pnas.0505266102)

67. Secco D, Wang C, Arpat BA, Wang Z, Poirier Y, Tyerman SD, Wu P, Shou H, Whelan J. 2012 The emerging importance of the SPX domain-containing proteins in phosphate homeostasis. New Phytol. 193, 842–851. (doi:10.1111/j.1469-8137.2011.04002.x)

68. Shi J, Hu H, Zhang K, Zhang W, Yu Y, Wu Z, Wu P. 2014 The paralogous SPX3 and SPX5 genes redundantly modulate Pi homeostasis in rice. J. Exp. Bot. 65, 859–870. (doi:10.1093/jxb/ert424)

69. Kiba T et al. 2018 Repression of nitrogen starvation responses by members of the arabidopsis GARP-type transcription factor NIGT1/HRS1 subfamily. Plant Cell 30, 925–945. (doi:10.1105/tpc.17.00810)

70. Müller F, Frentzen M. 2001 Phosphatidylglycerophosphate synthases from Arabidopsis thaliana. FEBS Lett. 509, 298–302. (doi:10.1016/S0014-5793(01)03163-5)

71. Jouhet J, Maréchal E, Bligny R, Joyard J, Block MA. 2003 Transient increase of phosphatidylcholine in plant cells in response to phosphate deprivation. FEBS Lett. 544, 63–68. (doi:10.1016/S0014-5793(03)00477-0)

72. Nakamura Y. 2013 Phosphate starvation and membrane lipid remodeling in seed plants. Prog. Lipid Res. 52, 43–50. (doi:10.1016/j.plipres.2012.07.002)

73. Salmela MJ, McMinn RL, Guadagno CR, Ewers BE, Weinig C. 2018 Circadian Rhythms and Reproductive Phenology Covary in a Natural Plant Population. J. Biol. Rhythms 33, 245–254. (doi:10.1177/0748730418764525)

74. Yerushalmi S, Green RM. 2009 Evidence for the adaptive significance of circadian rhythms. Ecol. Lett. 12, 970–981. (doi:10.1111/j.1461-0248.2009.01343.x)

75. West AC, Bechtold DA. 2015 The cost of circadian desynchrony: Evidence, insights and open questions. BioEssays 37, 777–788. (doi:10.1002/bies.201400173)

76. Green RM, Tingay S, Wang ZY, Tobin EM. 2002 Circadian rhythms confer a higher level of fitness to Arabidopsis plants. Plant Physiol. 129, 576–584. (doi:10.1104/pp.004374)

77. Michael TP, Salomé PA, Yu HJ, Spencer TR, Sharp EL, McPeek MA, Alonso JM, Ecker JR, McClung CR. 2003 Enhanced Fitness Conferred by Naturally Occurring Variation in the Circadian Clock. Science. 302, 1049–1053. (doi:10.1126/science.1082971)

78. Carré IA. 2002 ELF3: A circadian safeguard to buffer effects of light. Trends Plant Sci. 7, 4–6. (doi:10.1016/S1360-1385(01)02184-7)

79. Sakuraba Y, Bülbül S, Piao W, Choi G, Paek NC. 2017 Arabidopsis EARLY FLOWERING3 increases salt tolerance by suppressing salt stress response pathways. Plant J. 92, 1106–1120. (doi:10.1111/tpj.13747)

80. Chauhan BS, Johnson DE. 2008 Influence of Environmental Factors on Seed Germination and Seedling Emergence of Eclipta (Eclipta prostrata) in a Tropical Environment. Weed Sci. 56, 383–388. (doi:10.1614/ws-07-154.1)

81. Taylor DI, Nixon SW, Granger SL, Buckley BA, McMahon JP, Lin HJ. 1995 Responses of coastal lagoon plant communities to different forms of nutrient enrichment-a mesocosm experiment. Aquat. Bot. 52, 19–34. (doi:10.1016/0304-3770(95)00485-I)

82. Catford JA, Jansson R, Nilsson C. 2009 Reducing redundancy in invasion ecology by integrating hypotheses into a single theoretical framework. Divers. Distrib. 15, 22–40. (doi:10.1111/j.1472-4642.2008.00521.x)

83. Ericsson T. 1995 Growth and shoot: root ratio of seedlings in relation to nutrient availability. In Nutrient Uptake and Cycling in Forest Ecosystems, pp. 205–214. Springer Netherlands. (doi:10.1007/978-94-011-0455-5_23)

84. Olff H, Huisman J, Tooren BF Van. 1993 Species Dynamics and Nutrient Accumulation During Early Primary Succession in Coastal Sand Dunes. J. Ecol. 81, 693. (doi:10.2307/2261667)

85. Footitt S, Walley PG, Lynn JR, Hambidge AJ, Penfield S, Finch-Savage WE. 2020 Trait analysis reveals DOG1 determines initial depth of seed dormancy, but not changes during dormancy cycling that result in seedling emergence timing. New Phytol. 225, 2035–2047. (doi:10.1111/nph.16081)

