## Supplementary figures and images for "Examining the molecular mechanisms contributing to the success of an invasive species across different ecosystems"

### Figure S1

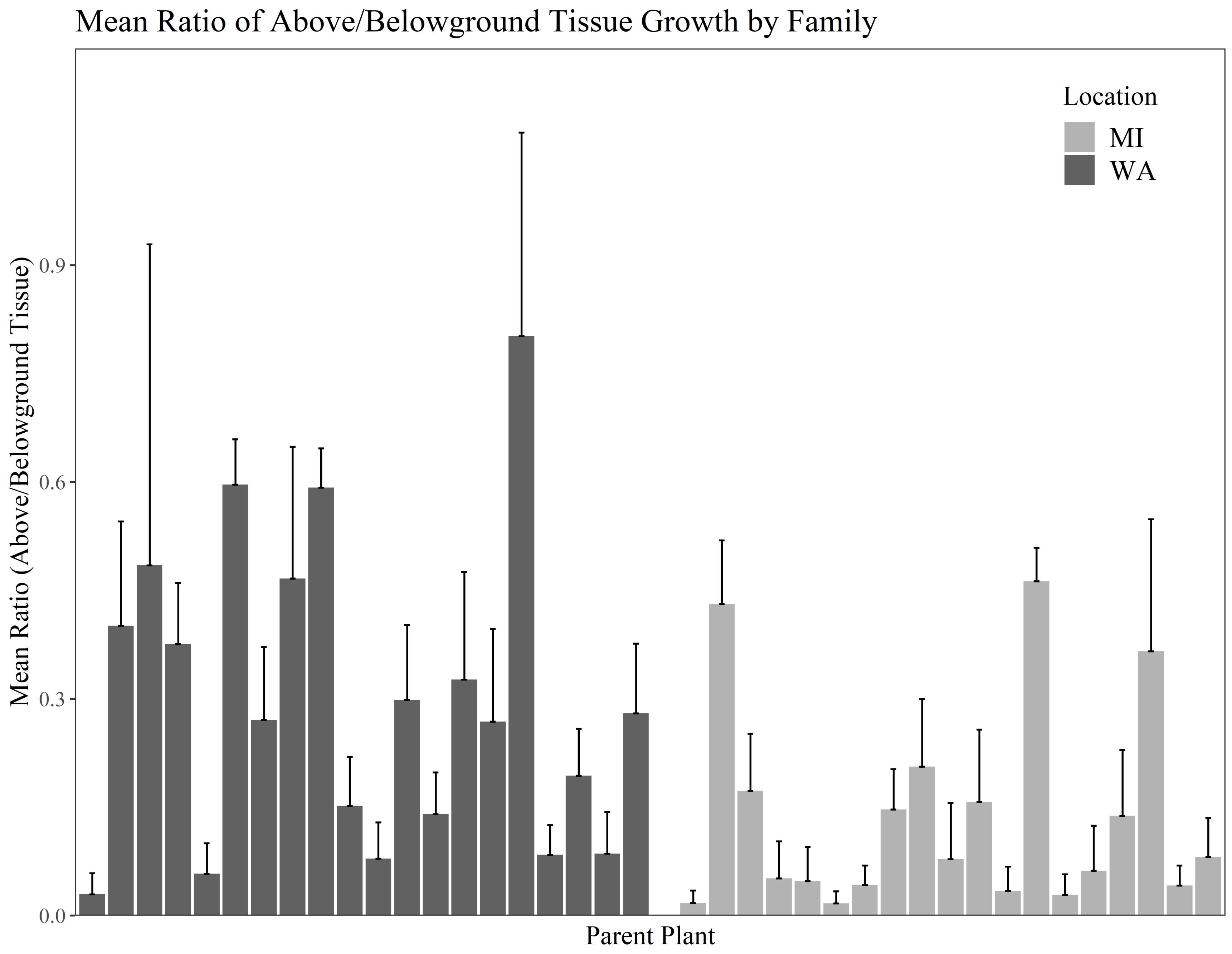
